# ROSE: a deep learning based framework for predicting ribosome stalling

**DOI:** 10.1101/067108

**Authors:** Sai Zhang, Hailin Hu, Jingtian Zhou, Xuan He, Tao Jiang, Jianyang Zeng

## Abstract

We present a deep learning based framework, called ROSE, to accurately predict ribosome stalling events in translation elongation from coding sequences based on high-throughput ribosome profiling data. Our validation results demonstrate the superior performance of ROSE over conventional prediction models. ROSE provides an effective index to estimate the likelihood of translational pausing at codon resolution and understand diverse putative regulatory factors of ribosome stalling. Also, the ribosome stalling landscape computed by ROSE can recover the functional interplay between ribosome stalling and cotranslational events in protein biogenesis, including protein targeting by the signal recognition particle (SRP) and protein secondary structure formation.

## 1 Background

The translation process, including translation initiation, elongation and termination, is a fundamental biological process that delivers genetic information to functional proteins in living cells. The dysregulation of translation has been shown to be associated with a variety of diseases, such as neurological disorder and cancers [1]. Elongation is a crucial step of mRNA translation after initiation, in which the ribosome scans the mRNA sequence and gradually grows the nascent peptide chain by appending new amino acids (Supplementary Fig. 1). Although numerous studies have shown that the local elongation rate along an mRNA sequence varies a lot, the underlying regulatory mechanisms of this phenomenon still remains unclear [2–6]. On the other hand, translation elongation plays essential roles in diverse aspects of protein biogenesis, such as differential expression, cotranslational folding, covalent modification and secretion [3, 4, 6]. In addition, the connection between the local elongation rate and human health is increasingly emerging, which further underscores the necessity of a good understanding on the regulatory mechanisms and functions of elongation dynamics [3, 7].

In recent years, ribosome profiling has emerged as a high-throughput sequencing-based approach to measure the ribosome occupancy on mRNAs at a translatome-wide level *in vivo* [2, 4, 5, 8, 9]. With an accurate inference of the ribosome A-site (i.e., the entry position of aminoacyl-tRNA) in a ribosome-protected fragment (also referred to as the ribosome footprint, ~30 nucleotides), ribosome profiling provides a genome-wide snapshot of translation elongation dynamics and offers a new angle to estimate translation efficiency. Currently, ribosome profiling has been widely used to study a number of important biological problems related to translation [2, 4, 5], such as the identification of novel or alternatively translated genome regions [10–17], and the discovery of critical regulatory factors in translational control and protein biogenesis [18–34]. Based on the current available large-scale studies involving ribosome profiling experiments, several databases, e.g., GWIPS-viz [35] and RPFdb [36], have been established to store these profiling data.

Although a large amount of sequencing data have been produced by ribosome profiling, researchers are still challenged by the complexity, heterogeneity and insufficient coverage of these data during the data analysis process [2, 4, 5, 37, 38]. Recently, deep learning has become one of the most popular and powerful techniques in the machine learning field [39, 40]. Its superiority over traditional machine learning models has been demonstrated in a wide range of applications, such as speech recognition [41], image classification [40] and natural language processing [42]. Specifically, deep learning has also been successfully applied to analyze large-scale genomic data and uncover notable biological patterns [43–50], such as the predictions of protein-nucleotide binding [43, 44, 50] and effects of noncoding sequence variants [47, 49]. In this work, we propose a deep learning based framework, called ROSE (RibosOme Stalling Estimator), to address the aforementioned challenges and model translation elongation dynamics based on the high-throughput ribosome profiling data.

In ribosome profiling experiments, ribosome stalling events can be inferred from ribosome footprint density and have been widely believed to negatively correlate with the local elongation rates [2, 4, 5]. Our framework ROSE casts the ribosome stalling modeling problem into a classification task, and predicts ribosome stalling using a deep convolutional neural network (CNN) with encoded sequence features. ROSE is trained through a supervised manner based on both human and yeast ribosome profiling data to revisit evolutionarily conserved observations about ribosome stalling. Our validation results demonstrate that ROSE can greatly outperform conventional machine learning methods for predicting ribosome stalling. We further show that the predictions of ROSE generally correlate with diverse putative regulatory factors of ribosome stalling, including codon usage bias, tRNA adaptation, codon cooccurrence bias, pro-line codons, N^6^-methyladenosine (m^6^A) modification, mRNA secondary structure and protein-nucleotide binding, and provide a useful index to characterize the likelihood of ribosome stalling at codon resolution. Moreover, our comprehensive *in silico* studies reexamine several regulatory relations between elongation dynamics and cotranslational events in protein biogenesis. In particular, our analysis confirms an interesting interplay between elongation dynamics and the signal recognition particle (SRP) binding of transmembrane (TM) segments. Our results suggest that ribosomes tend to stall with high probability at a position ~50 codons downstream from a TM segment, which may promote the molecular recognition by SRP. In addition, our studies show that protein secondary structure elements (SSEs) and their transition patterns tend to be highly correlated with the the likelihood of ribosome stalling, which implies an essential regulatory effect of elongation dynamics on cotranslational folding. Furthermore, our intergenic analysis suggests that the enriched ribosome stalling events at the 5’ ends of coding sequences may be functionally important in the modulation of translation efficiency. These results demonstrate that ROSE can offer an effective tool for analyzing large-scale ribosome profiling data and provide new perspectives on understanding the landscape of ribosome stalling.

## 2 Results

### 2.1 Designing and training ROSE

We propose a deep learning based framework, called RibosOme Stalling Estimator (ROSE), to analyze large-scale ribosome profiling data and study the contextual regulation of ribosome stalling and its potential functions in protein biogenesis (Fig. 1a). Unlike previous work that characterized translation elongation dynamics using the stochastic simulation approaches [20, 25, 26] and density estimation [51, 52], ROSE formalizes the modeling problem as a classification task, in which the resulting prediction score can be used to measure the probability of a ribosome stalling event. In this classification framework, codon positions with normalized ribosome footprint densities beyond two standard deviations of the density distribution are defined as positive samples (foreground), which represent the occurrences of ribosome stalling, while the remaining sites are regarded as negative samples (background; Supplementary Fig. 2). This threshold is selected to best correlate the normalized reads with the model predictions in a separate validation dataset (Supplementary Fig. 3 and Methods).

**Figure 1:**
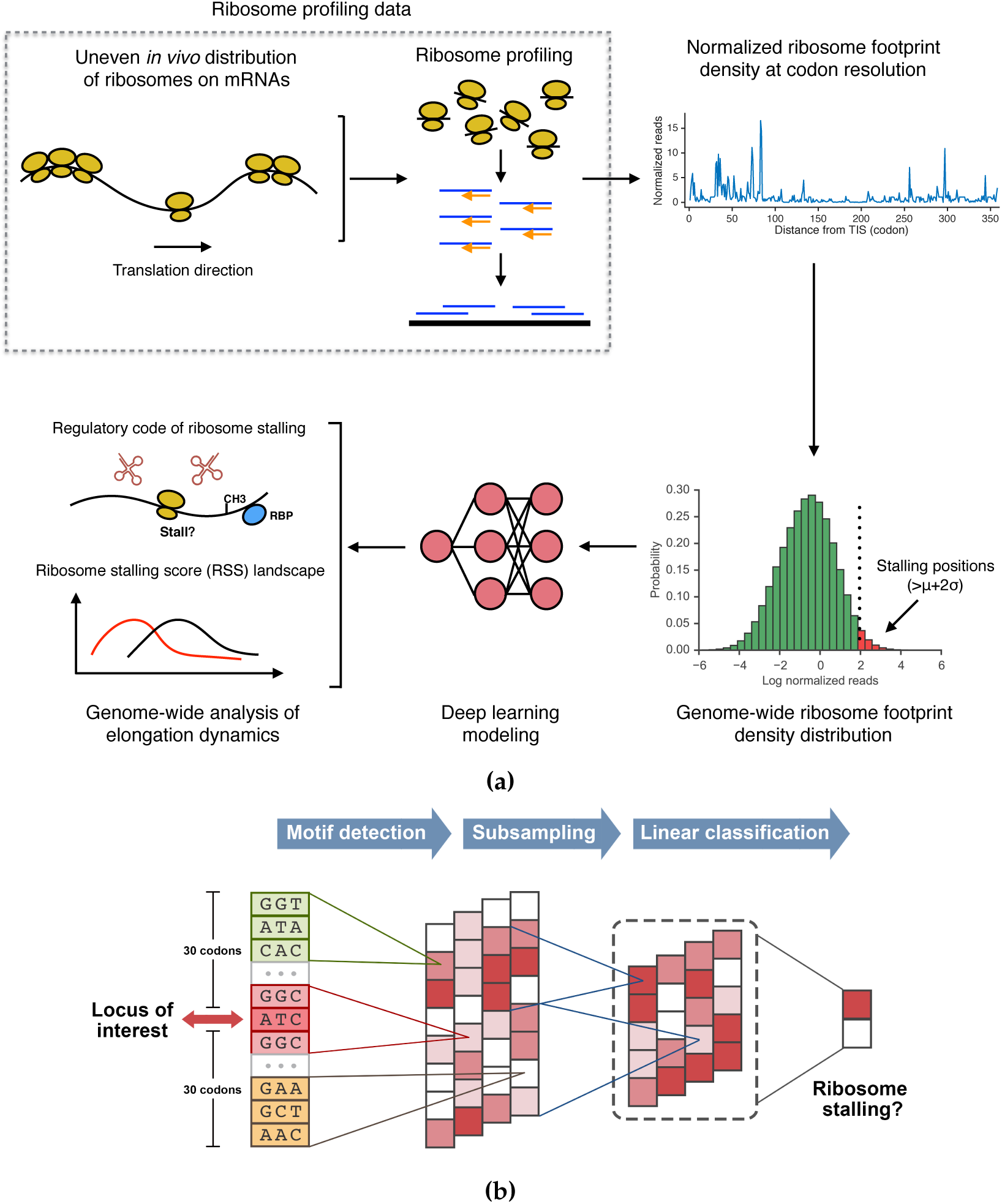
The ROSE pipeline and the CNN model. (a) Schematic overview of the ROSE pipeline. The codon sites with normalized ribosome footprint densities beyond two standard deviations are regarded as positive samples, which represent the ribosome stalling positions, to train a deep CNN model. Then the sequence profiles of individual codon sites along the genome are fed into the trained CNN to compute the distribution of ribosome stalling, which can be further used to study the potential factors affecting ribosome stalling and analyze the genome-wide landscape of translation elongation dynamics. (b) Schematic illustration of the CNN model used in the ROSE pipeline. More details can be found in the main text.

We assume that a ribosome stalling event is primarily determined by its surrounding sequence. The codon position of interest, i.e., the ribosome A-site, is first extended both upward and downward by 30 codons, which yields the codon sequence profile of a putative stalling event. We then encode this sequence and feed it into a deep convolutional neural network (CNN) to learn the complex relations between ribosome stalling and its contextual features (Fig. 1b and Methods). We call the prediction score directly output by the CNN the *intergenic ribosome stalling score*, which is also termed interRSS (Methods). The name “interRSS” comes from the fact that all the scores along the genome are calculated by a universal model and can be compared intergenetically/globally under the same criterion. To further eliminate the possible bias among different genes and facilitate the study on the interplay between intragenic/locally factors (e.g., the binding of the signal recognition particle (SRP) on transmembrane segments) and elongation dynamics, we also normalize interRSS within each gene and obtain a local index, called the *intragenic ribosome stalling score*, which is also termed intraRSS (Methods). Here we basicly follow the same terminologies “intergenic” and “intragenic” from [6]. We collectively call interRSS and intraRSS the *ribosome stalling score* (RSS). In principle, RSS can be considered as an estimate of the likelihood of ribosome stalling. A higher RSS generally indicates a higher probability of ribosome stalling at the corresponding codon position and vice versa.

ROSE relies on a number of motif detectors (i.e., convolution operators) to scan the input sequence and integrate those stalling relevant motifs to capture the intrinsic contextual features of ribosome stalling (Fig. 1b and Methods). Unlike previous CNN architectures used for analyzing biological data [44, 47], our new CNN framework includes multiple *parallel* convolution-pooling modules, which can not only significantly reduce the model complexity, but also alleviate the potential overfitting problem (Methods). The standard error back-propagation algorithm is used to learn the network parameters of the CNN model [53]. We also deploy several optimization techniques, including *L*_2_-regularization [54], dropout [54, 55] and early stopping [54], to further overcome the overfitting problem (Methods).

To alleviate the need for tuning model hyperparameters depending on expert knowledge and optimize the model settings, here we propose an efficient strategy for large-scale automatic model selection, i.e., calibrating the model hyperparameters without any human intervension (Methods). To further boost the prediction performance, we also implement an ensemble version of ROSE (termed eROSE), in which 64 CNNs are initialized and trained independently, and then the average result is used as the final prediction score (Methods).

### 2.2 ROSE accurately predicts ribosome stalling

In this study, we mainly focused on eukaryotic cells, including human and yeast cells. We first used two datasets downloaded from GWIPS-viz [35], including a human dataset of lymphoblastoid cell lines (LCLs) (denoted by Battle15) [56] and a yeast dataset of *S. cerevisiae* (denoted by Pop14) [25] to train our deep learning model and evaluate its prediction performance. More details about data preparation, preprocessing and normalization can be found in Methods.

We first compared the performance of ROSE to that of a conventional prediction model, called gkm-SVM, which is developed mainly based on the support vector machine (SVM) and has been successfully applied in various analyses of genomic sequence data [57, 58]. Our tests on both human and yeast datasets showed that ROSE greatly outperformed gkm-SVM with an increase in the area under the receiver operating characteristic curve (AUROC) by up to 18.4% (Figs. 2a and 2b). In particular, the ensemble version of ROSE (also termed eROSE) consistently had superior performance compared to the single version (also termed sROSE). To validate the effectiveness of our parallel CNN architecture, we also implemented three sequential architectures that stacked two convolution-pooling modules with different kernel sizes in the convolutional layers before the output layer, and found that sROSE greatly outperformed those sequential CNNs (Fig. 2c).

**Figure 2:**
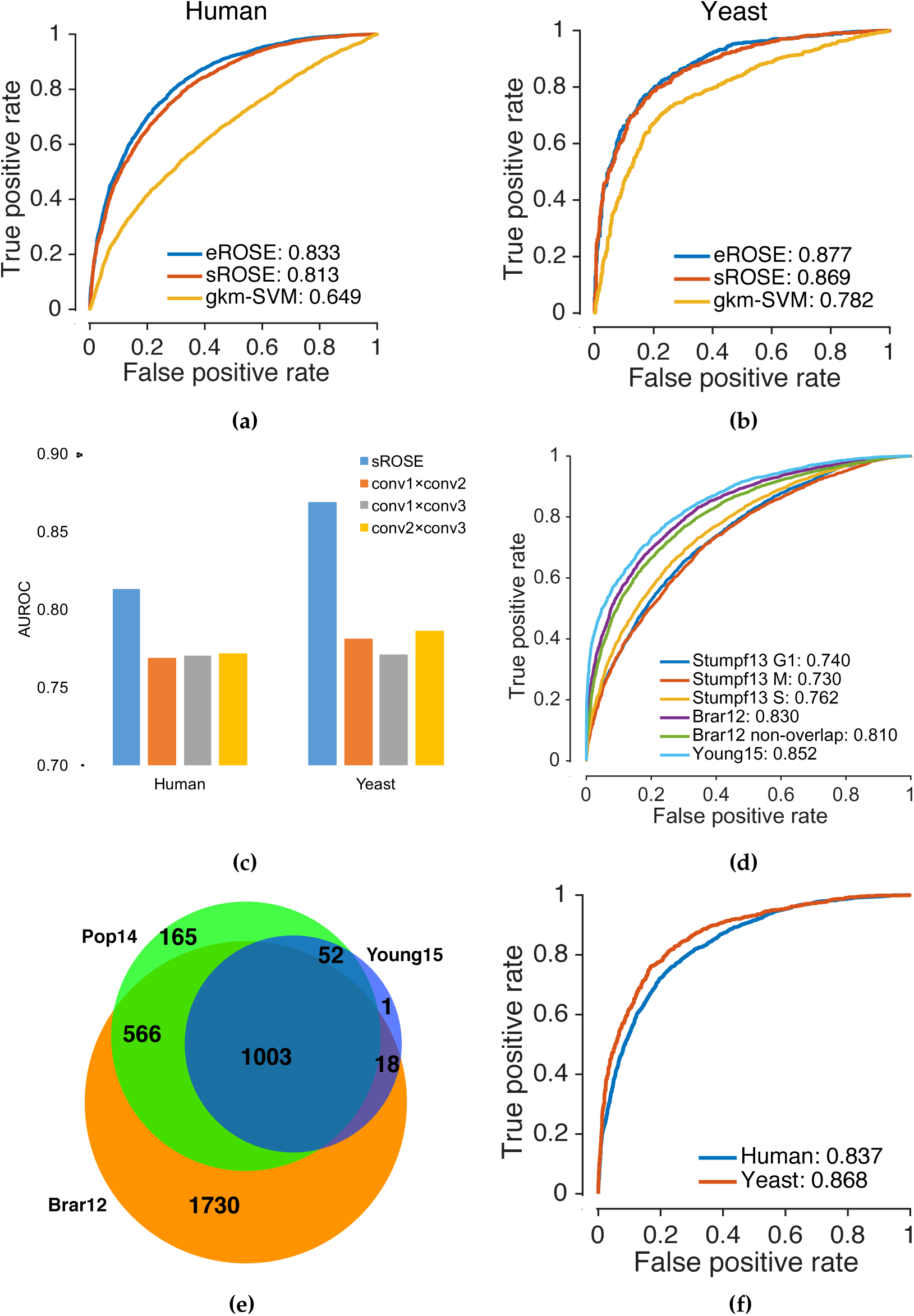
Performance evaluation of ROSE. (a) and (b) The receiver operating characteristic (ROC) curves and the area under the corresponding ROC curve (AUROC) scores on the human (Battle15) and yeast (Pop14) test datasets, respectively. (c) Comparison of AUROCs between parallel and sequential CNN architectures. “sROSE” and “eROSE” stand for the ROSE frameworks with one (single) and 64 (ensemble) CNNs, respectively. conv1, conv2 and conv3 represent the convolutional-pooling modules with kernel sizes of the convolutional layers corresponding to short, medium and long ranges used, respectively. “×” denotes the stacking operation in the sequential architecture. See the main text for more details. (d) The ROC curves and the corresponding AUROC scores of the cross-study tests on additional human (Stumpf13 G1, M and S) and yeast (Brar12 and Young15) datasets, respectively. The Brar12 non-overlapping dataset includes 1,748 genes with sufficient (over 60%) ribosome profiling coverage in the Brar12 dataset but in not the Pop14 dataset. (e) The Venn diagram of the three yeast datasets regarding sufficiently covered genes. (f) The ROC curves and the corresponding AUROC scores on the ramp regions.

We further performed multiple cross-study analyses to examine the generalizability of ROSE over other ribosome profiling datasets with different experimental conditions, e.g., cell lines/strains and cycloheximide treatment (Methods). In particular, we used five additional datasets for cross-study validation, including three human datasets from different cell cycle stages (i.e., G1, S and M phases) of Hela cells [59] (denoted by Stumpf13 G1, S and M, respectively) and two other yeast datasets of strain SK1 [60] and BY4741 [61] (denoted by Brar12 and Young15, respectively). Here, the ROC curves were obtained by applying the trained eROSE of the corresponding species to these validation data. As expected, these cross-study validations displayed robust prediction performance of ROSE with only a moderate decrease in AUROC scores for both human and yeast (Fig. 2d). Interestingly, we observed less variance of prediction performance for yeast, possibly due to a less sophisticated regulation mechanism of translation elongation dynamics in yeast. Moreover, we noted there were a number of genes with sufficient ribosome profiling coverage (>60%) in Brar12 dataset but not in Pop14 dataset (Fig. 2e). Notably, our model can still recover the ribosome stalling events in these non-overlapped genes, with the prediction accuracy comparable to that of the whole Brar12 dataset (Fig. 2d), illustrating the ability of our model to successfully identify ribosome stalling events in regions with low sequencing coverage in ribosome profiling data. These results indicated that although trained only based on a single dataset for each specific organism, ROSE can still capture intrinsic sequence preferences in a ribosome stalling event, which are more or less conserved in different cell lines/strains and experimental conditions.

It has been widely observed that the first 30-50 codons of a coding sequence (CDS) are often enriched with rare codons, and create a “ramp” to reduce the elongation rate during the initial translation elongation process [62, 63]. Such a ramp sequence has been proposed to be universal in prokaryotic and eukaryotic genes, and has been suggested to serve the purpose of reducing the likelihood of downstream ribosomal traffic jams and thus increasing the overall translation efficiency [6, 62, 63]. To focus solely on the elongation process and remove the possible biases introduced by the ramp regions, here we excluded all the reads of these regions, i.e., the first 50 codons at the 5’ ends of coding sequences, from our training data. On the other hand, we showed that even without using any training sample from ramp regions, ROSE can still successfully predict ribosome stalling in these regions, with the AUROC scores above 83.0% (Fig. 2f). Since ramp sequences are of particular interest in the literature, we further carried out several functional analyses of ribosome stalling in these regions, and found that the enriched ribosome stalling events in ramp regions may be involved in modulating the translation efficiency of important genes at the elongation level (Supplementary Fig. 4). More details can be found in Supplementary Notes.

### 2.3 ROSE associates ribosome stalling with putative regulatory factors

Next, we analyzed several putative factors that may play important roles in regulating ribosome stalling during translation elongation. Previous studies [4, 8, 10, 19, 20, 22, 25–27, 64, 65] have provided various mechanistic insights into translation elongation dynamics based on statistical analysis. With stringent normalization procedures as well as superior prediction performance, ROSE enables one to systematically investigate diverse factors that may associate with ribosome stalling (Methods). Here, we mainly focused on codon usage bias, codon cooccurrence bias, pro-line codons and mRNA N^6^-methyladenosine modification, and studied how they correlate with ribosome stalling and elongation dynamics.

#### Codon usage bias

Systematic variation has been observed in codon usage across species, among genes in a genome, or even within a gene [66]. Such a codon usage bias has been demonstrated to be crucial in various cellular functions, such as splicing control, maintenance of translation fidelity and regulation of protein folding [3, 7]. Particularly, rare codons are normally highly associated with low translation elongation rates and high ribosome stalling potential, which has been widely believed to be responsible for proper protein folding [3, 6, 7, 66, 67]. Several metrics have been proposed to measure the codon usage frequency, such as the codon adaptation index (cAI) [68] and the %MinMax score [69]. Here, we used ROSE to reexamine how the codon rareness correlates with ribosome stalling based on both cAI and %MinMax profiles. Specifically, we scanned the ribosome occupancy sites along a genome, and compared the intraRSSes of those sites enriched with rare codons to those of the background (Methods). We calculated cAI for both ribosome A-and P-sites, and %MinMax for the local region around the ribosome A-site (i.e., five codons both upstream and downstream from the A-site). Consistent results were observed for both human and yeast, i.e., those sites enriched with rare codons displayed significantly higher intraRSSes than the background (Figs. 3a, 3b and Supplementary Figs. 5a, 5b; *P* < 10^−65^ for cAI and *P* < 10^−25^ for %MinMax, one-sided Wilcoxon rank-sum test). These results implied that the codon usage bias may be an important factor to modulate ribosome stalling.

**Figure 3:**
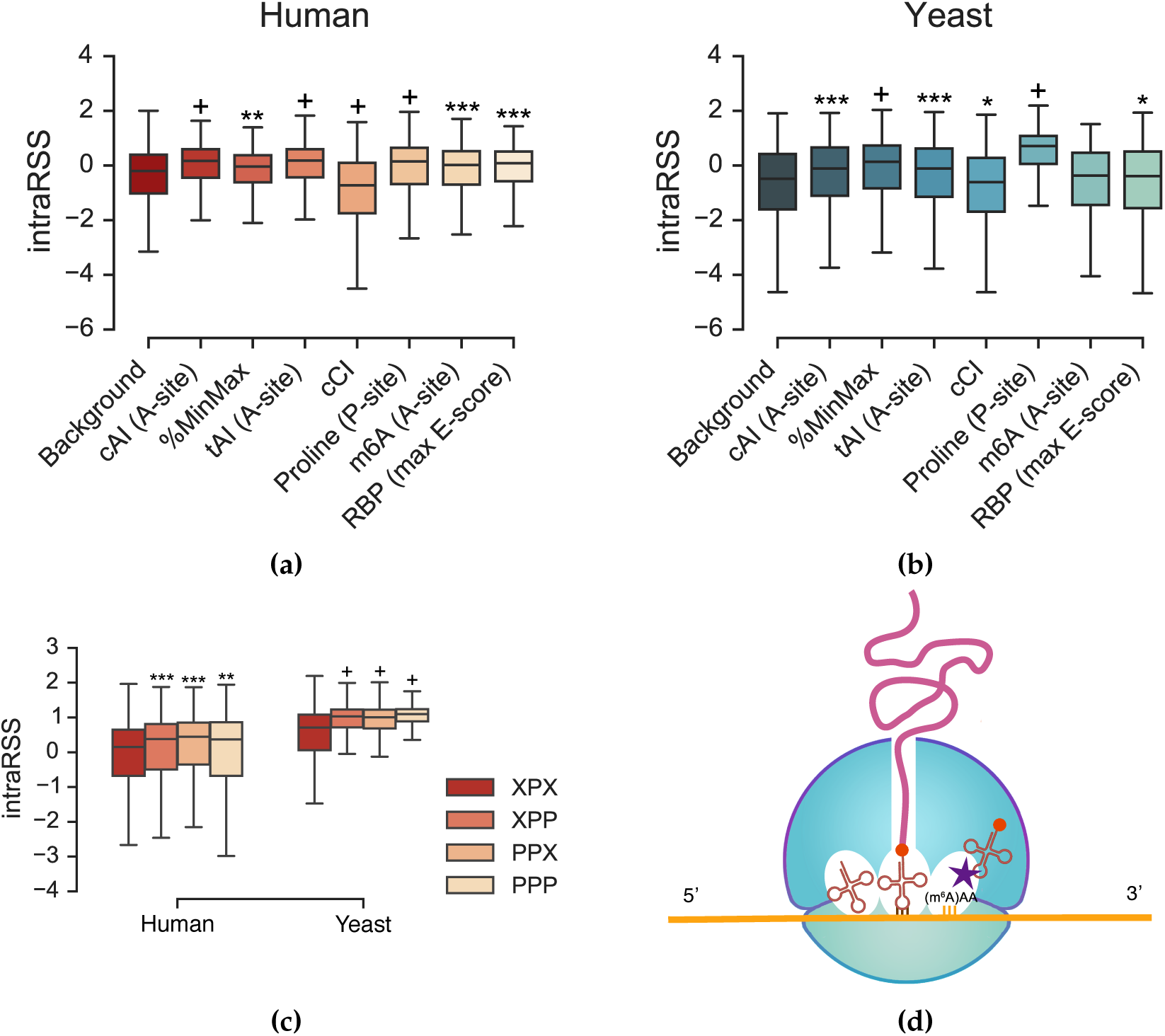
A comprehensive reexamination on the relations between diverse putative regulatory factors and ribosome stalling using ROSE. (a) and (b) The comparisons of intraRSS between the codon sites enriched with individual factors and the background for human and yeast, respectively. (c) The comparisons of intraRSS between the single-peptide pattern of proline (i.e., XPX) and the multiple-peptide patterns of proline, including dipeptide (i.e., XPP and PPX) and tripeptide (i.e., PPP), where “P” and “X” stand for proline and any non-proline amino acid, respectively. (d) A schematic illustration of the m^6^A modification of a codon (e.g., AAA) to delay tRNA accommodation during translation elongation. *: 5 × 10^−25^ < *P <* 1 × 10^−2^; **: 5 × 10^−50^ < *P* ≤ 5 × 10^−25^; ***: 5 × 10^−100^ < *P* ≤ 5 × 10^−50^; +: *P* ≤ 5 × 10^−100^; one-sided Wilcoxon rank-sum test.

#### Codon cooccurrence bias

The codon cooccurrence bias, i.e., the non-uniform distribution of synonymous codon orders, can affect translation elongation dynamics [6, 70]. Previous studies have suggested that after being recharged, tRNAs may still stay around the ribosome, and tRNA recycling can modulate the translation elongation rate [6, 70]. Under this hypothesis, we would expect the highly isoaccepting-codon-reused regions to be depleted of ribosome stalling events. To examine this problem, we first defined a new metric, called the *codon cooccurrence index* (cCI), which measures the autocorrelation (i.e., reuseness) of isoaccepting codons in a local region. Precisely speaking, given the codon of interest at position *i*, we only considered its local region [*i − w, i* + *w*], where *w* stands for the window size. For each codon at position *p* ∈ [*i – w, i* + *w*], we checked whether it had an isoaccepting codon in the upstream region [*i − u, p* − 1]. We used notation iso_*p*_ to represent this indicator, that is, iso_*p*_ = 1 if the indicator holds true, and iso_*p*_ = 0 otherwise. Thus, the cCI at position *i* was defined as

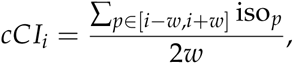
in which we set *w* = 5 and *u* = 30. Our studies on both human and yeast datasets showed that those regions enriched with autocorrelated codons (i.e., with high cCI scores) displayed significantly lower intraRSSes than the background (Figs. 3a and 3b; *P* < 10^−9^ by one-sided Wilcoxon rank-sum test). This result provided a novel evidence to support the argument that the codon cooccurrence bias may be related to the regulation of ribosome stalling.

#### Proline codons

The unique structure of proline side chain is generally associated with a relatively low efficiency in its peptide bond formation, which may slow down translation elongation [19, 67, 71–76]. Several studies have confirmed the relatively low translation elongation rates at proline codons [19, 67]. Here we performed an extended study on the relation between proline codons and ribosome stalling using ROSE. In particular, four peptide patterns of proline were investigated, including XPX, XPP, PPX and PPP, in which the three positions correspond to the ribosome E-, P-and A-sites, respectively, and “P” and “X” represent proline and non-proline amino acids, respectively. The comparative analysis on both human and yeast datasets showed that these peptide patterns of proline displayed significantly higher intraRSSes than the background (Figs. 3a and 3b; *P* < 10^−100^ by one-sided Wilcoxon rank-sum test). In addition, the dipeptide and tripeptide patterns of proline, including XPP, PPX and PPP, showed significantly higher intraRSSes than XPX (Fig. 3c; *P* < 10^−40^ by one-sided Wilcoxon rank-sum test). This result implied that a codon sequence with more prolines may yield a higher chance of ribosome stalling.

#### N^6^-methyladenosine modification

Notably, we found that the mRNA modification N^6^-methyladenosine (m^6^A) within codons can also correlate with ribosome stalling (Fig. 3d). N^6^-methyladenosine is probably the most prevalent post-transcriptional modification in mRNAs and plays vital roles in regulating mRNA stability and translation efficiency [77,78]. Most m^6^A sites are distributed in mRNA regulatory regions such as 3’-UTRs and the locations around stop codons, where the m^6^A “readers” such as YTHDF1 and YTHDF2 can bind to dynamically regulate gene expression [77]. Moreover, it has been reported that m^6^A in the CDS can modulate the translation initiation to promote translation efficiency [78]. During translation, the additional methyl group of an m^6^A site may act as a stumbling block to ribosomes and thus result in a translational pause. Recently, Choi *et al.* elucidated that the m^6^A-modified codons at the ribosome A-sites can reduce the translation elongation rate in *E.coli* [79]. Thus, it is reasonable to hypothesize that the m^6^A marks in eukaryotic cells may also be closely correlated with the ribosome stalling tendency. We used ROSE to test this hypothesis based on the translatome-wide m^6^A mapping obtained from the known single-nucleotide resolution sequencing data, including two human datasets (denoted by Linder15 [80] and Ke15 [81], respectively) and one yeast dataset (denoted by Schwartz13 [82]). The analysis results showed that in human, those codons modified by m^6^A at the ribosome A-sites had a significantly higher ribosome stalling tendency than the background (Fig. 3a and Supplementary Fig. 5c; *P* = 9.61 × 10^−13^ for Linder15 and *P* = 2.20 × 10^−64^ for Ke15, one-sided Wilcoxon rank-sum test). To ensure that such an increase of intraRSS did not result from the underlying adenine nucleotides in the codon sites of interest, we also constructed a control dataset which contained 10, 000 randomly-selected codon sites covering the adenine nucleotides but without m^6^A modification. In the control test, those m^6^A codon sites from the human translatome still exhibited higher intraRSSes than the control dataset (Supplementary Fig. 5d; *P* = 1.76 × 10^−9^ for Linder15 and *P* = 4.52 × 10^−54^ for Ke15, one-sided Wilcoxon rank-sum test). For yeast, the difference of intraRSS between the m^6^A codon sites and the background was insignificant (Fig. 3b and Supplementary Fig. 5d; *P* > 0.05 by two-sided Wilcoxon rank-sum test). This result may be due to the fact that the number of codons modified by m^6^A in the yeast dataset was quite limited (only 278 samples after read mapping), and the m^6^A modification was only observed in yeast meiosis [82]. Overall, our analysis suggested a novel relation between the m^6^A modification and translation elongation in human and yeast, which may further expand the previous conclusion drawn from the *E. coli* data [79].

#### Other factors

We also applied ROSE to analyze the relations between ribosome stalling and other factors, including tRNA adaptation, mRNA secondary structure and protein-nucleotide binding, and found that they may also associate with ribosome stalling (Figs. 3a and 3b and Supplementary Figs. 5 and 6). On the other hand, different from several previous study results [25, 64, 83–85], we did not observe a strong association between positively-charged amino acids and ribosome stalling according to our prediction results (Supplementary Fig. 7). More details can be found in Supplementary Notes.

### 2.4 ROSE captures the landscape of ribosome stalling

As the RSS output by ROSE correlates with diverse putative factors that may affect ribosome stalling, it can be regarded as an effective indicator of translation elongation dynamics. On the other hand, due to the uneven nature of elongation dynamics, each gene can have a specific intra-genic landscape of ribosome stalling, which may be important for many cellular functions, e.g., modulating the distribution of ribosomes along an mRNA [62, 63], regulating the protein cotranslational folding process [86], and assisting protein translocation across membranes [87]. Here, we analyzed the RSS landscape by relating it to several cotranslational events in protein biogenesis, including protein secondary structure formation and protein targeting by the signal recognition particle (SRP).

#### RSS correlates with protein secondary structure

Although codon bias has been shown to associate with protein secondary structure elements (SSEs) and cotranslational folding [3, 6, 87, 88], it lacks more direct evidence to verify that such an association is imparted from the local elongation dynamics. Here, we sought to probe the relations between the protein SSEs and ribosome stalling based on the RSS computed by ROSE. In particular, we first derived a set of non-redundant protein chains across human and yeast genomes from the Protein Data Bank (PDB) [89], in which BLAST [90] with the sequence-similarity cutoff *P* = 10^−7^ was used to compare two protein sequences. The SSEs of these protein chains (5,054 from human and 766 from yeast) were then determined based on the mapping to the DSSP database [91, 92], which contains the experimentally-determined secondary structure assignments for the protein sequences in the PDB. We then investigated intraRSS landscapes of different SSE patterns, including a single chain of alpha helix (H), beta stand (B) or random coil (C), and transitions between different SSEs.

To obtain the average position-specific intraRSSes of a certain SSE pattern, all the eligible SSE-aligned sequences with a particular window size were extracted from the genome with five flanking amino acids on both sides, and then the mean intraRSS of each position was calculated. Note that here we mainly considered the intraRSSes of those codons at the ribosome P-sites, where the corresponding amino acids are concatenated to the nascent peptides (Supplementary Fig. 1). Overall, we found that with the window size of six, all the tendencies of the intraRSS change for individual SSE patterns were species independent (Figs. 4a and 4b; Spearman correlation coefficient *R* > 0.6). To further eliminate the bias that may be caused by the window size, we also repeated the same analysis procedure with the window size of ten, and found that five out of seven SSE patterns still showed similar trends for both human and yeast (Supplementary Fig. 8; Spearman correlation coefficient *R* > 0.5). The remaining two SSE patterns displayed relatively weak agreements in intraRSS tendencies (Supplementary Fig. 8; Spearman correlation coefficient *R* < 0.1), which was probably attributed to the dramatic drop of sample size in both datasets (i.e., from thousands to tens).

**Figure 4:**
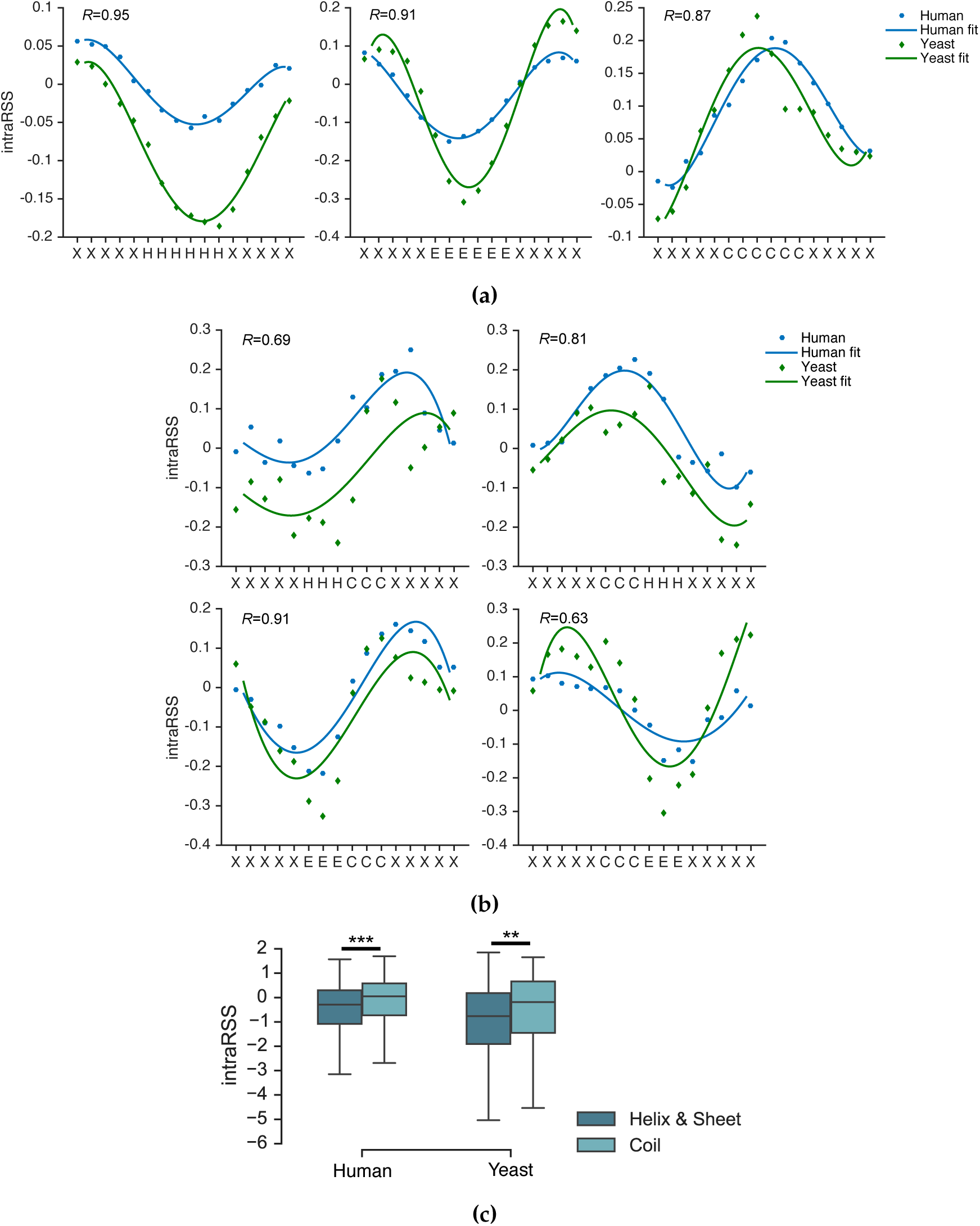
The intragenic RSS landscape reveals that ribosome stalling associates with protein secondary structure. (a) The intraRSS landscapes of alpha helix, beta strand and random coil regions. (b) The intraRSS landscapes of the SSE transition regions. “H”, “E” and “C” stand for alpha helix, beta strand and random coil, respectively, while “X” represents any SSE type in the flanking regions on both sides. Polynomial curve fitting of degree four was used to show the general intraRSS tendency. The Spearman correlation coefficients between human and yeast intraRSS tendencies were calculated. (c) The overall comparisons of intraRSS between the structured (i.e., alpha helix and beta strand) and random coil residues. **: 5 × 10^−50^ < *P ≤* 5 × 10^−25^; ***: 5 × 10^−100^ < *P* ≤ 5 × 10^−50^; one-sided Wilcoxon rank-sum test.

We further compared the intraRSSes of the structured (i.e., alpha helix or beta strand) and random coil residues at the ribosome P-sites. Consistent with the previous report that frequent codons were usually enriched in the structured regions while depleted in the random coils [87], our results showed a significantly higher stalling probability in the coils than in the alpha helix or beta strand regions (Fig. 4c; *P* < 10^−25^ by one-sided Wilcoxon rank-sum test). Furthermore, we examined the tendency of the intraRSS change along a protein secondary structure fragment. As expected, the intraRSS landscape showed a lower chance of stalling in the middle of a structured region while a higher chance in the middle of a coil region, when compared to the corresponding flanking regions on both sides (Fig. 4a and Supplementary Fig. 8a). This behavior was reminiscent of another previous study on the relations between codon frequency and protein secondary structure, in which the tRNA adaptation index (i.e., tAI) was mainly used as an indicator of the elongation rate [88]. Our intraRSS landscape showed a similar trend to the previous finding that the transitions from structured to coil regions generally accompanied an increase in the stalling probability on the transition boundaries (Fig. 4b and Supplementary Fig. 8b). In addition, the opposite transitions (i.e., from coil to structured regions) exhibited roughly symmetrical trends in the change of intraRSS (Fig. 4b and Supplementary Fig. 8b). Notably, the ribosome stalling positions revealed by intraRSS here were slightly different from those reported in [88]. In particular, our results displayed more symmetrical stalling positions than the previous results, suggesting that ribosomes are prone to stall at the coil residues near the transition boundaries. Admittedly, we cannot exclude the influence of other factors on these findings, such as the database upgrade or the discrepancy of SSE assignment between different databases (e.g., JOY [93] vs. DSSP [91, 92]). Nevertheless, our results indicated the necessity of incorporating other putative factors in addition to tRNA adaptation to better estimate the ribosome stalling tendency and elongation dynamics.

#### RSS associates with SRP recognition

Next, we investigated whether the RSS landscape can reflect the elongation process that regulates the coupling between the protein translation and translocation activities. We were particularly interested in the interplay between the translational pause and the signal recognition particle (SRP) binding of transmembrane (TM) segments (Fig. 5a). We expected that our model would effectively capture the ribosome stalling events encoded by the heterogeneity of amino acid composition in and around the TM domains.

**Figure 5:**
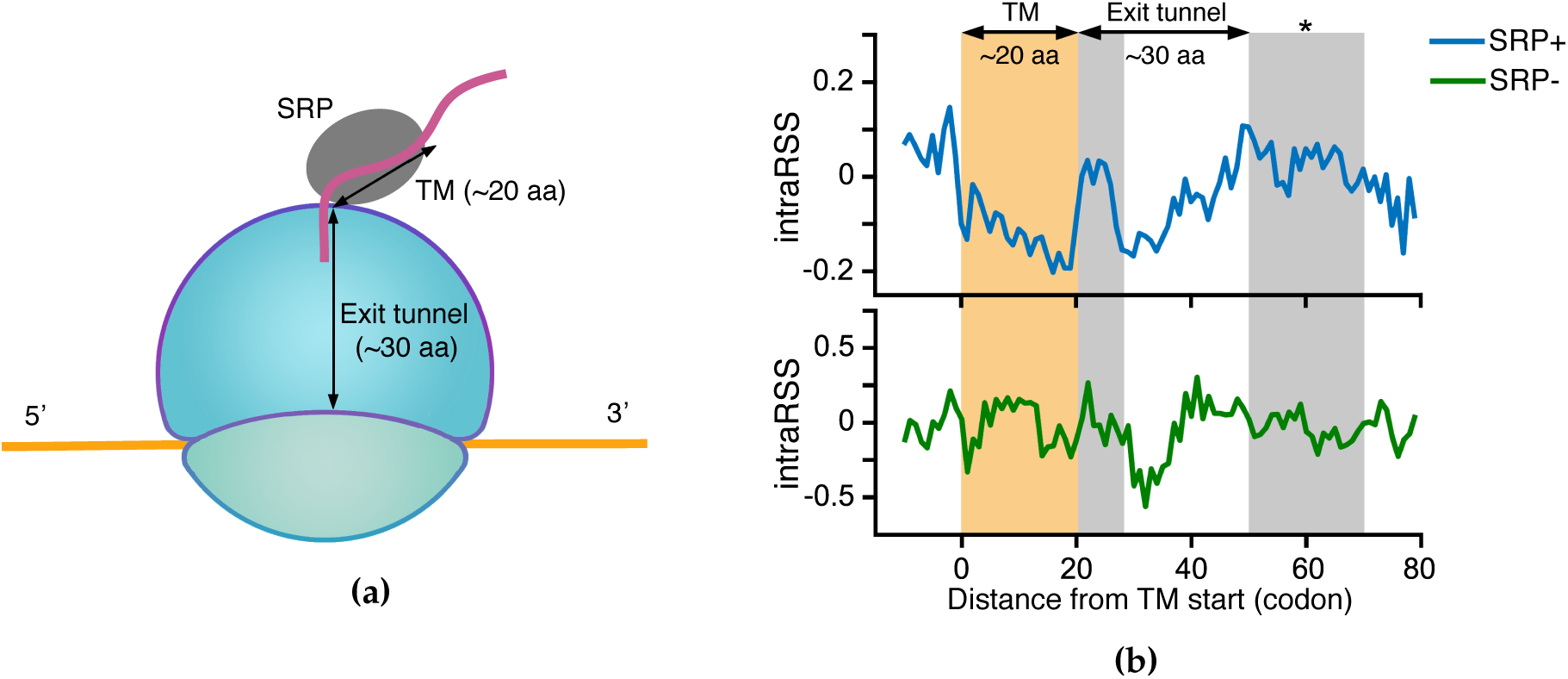
The intragenic RSS landscape shows that ribosome stalling correlates with the SRP binding of TM segments. (a) A schematic illustration of the SRP binding of a TM segment during translation elongation. (b) The comparison of intraRSS tendency between the TM segments with (SRP+) and without SRP binding (SRP-) in yeast, in which all the protein sequences were aligned with regard to the start of the TM segment whose position was indexed as zero. The yellow rectangle covers the TM segment, while the grey rectangles represent two intraRSS peaks downstream the TM segment. *: 5 × 10^−25^ < *P* < 1 × 10^−2^; one-sided Wilcoxon rank-sum test.

We first downloaded all the available TM protein sequences of human and yeast as well as the corresponding TM domain information from the Uniprot database [94]. To avoid the biases that may be caused by the influence between different TM segments, here we only considered the single-pass integral proteins and the last TM segments of the multispan TM proteins, which resulted in 4,235 human and 561 yeast proteins. For yeast proteins, we also excluded 65 TM sequences that are not bound by SRP according to the previous experimental study [95]. To characterize the intraRSS landscape along the elongation process, all the protein sequences were aligned with regard to the start of the TM segment whose position was indexed as zero, and then the mean intraRSS of each codon between positions −10 and +80 was calculated.

We first focused on the yeast TM proteins, whose translation has been previously characterized both computationally and experimentally [96]. The intraRSS landscape computed by ROSE captured two major stalling events during the TM protein translation process (Fig. 5b). The first stalling event after the TM start occurred right at the end of the TM segment, where the structured TM segment (majorly alpha helix) is transited to a more flexible intracellular region. This result agreed well with our previous conclusion about the relations between RSS and protein secondary structure (Fig. 4b and Supplementary Fig. 8b). The other intraRSS peak, spanning positions from +50 to +70, probably represented the intrinsic stalling to promote the nascent-chain recognition by SRP, which was consistent with the previous report [96]. Indeed, a TM segment generally contains ~20 residues and the length of the ribosome exit tunnel is ~30 residues. Thus position +50 is approximately the place where the translated TM segment emerges from the exit tunnel and is bound by SRP.

To further verify whether this intraRSS peak was truly relevant to SRP binding, we also examined the intraRSS landscape of the TM segments that were not associated with SRP binding (termed SRP-), which were obtained from the previous experimental study [95]. Compared to the above result on the sequences with SRP binding (termed SRP+), the first stalling event remained at position +20, while the intraRSS peak in the putative SRP-associated stalling region (i.e., positions from +50 to +70) was significantly diminished (Fig. 5b; *P* = 1.5 × 10^−3^ by one-sided Wilcoxon rank-sum test), which indicated that intraRSS is an excellent indicator of ribosome stalling that regulates the synthesis of a TM domain. Notably, the TM segments at positions between 0 and +20 of the SRP-proteins showed a remarkable increase in intraRSS compared to the result of the SRP+ proteins, suggesting an alternative membrane targeting may occur in the absence of SRP binding. To probe whether our position alignment scheme can induce bias in our analysis, we also performed a similar study in which the TM ends were aligned together, from which we drew the same conclusion (Supplementary Fig. 9a).

For human, as we lacked the SRP recognition data, it was difficult to separate the TM domains bound and unbound by SRP. Here, we only analyzed the intraRSS landscape of a mixed human dataset that did not distinguish the TM segments with and without SRP binding. In this analysis, although we still observed a similar intraRSS peak near the end of the TM segment, the peak signal around position +50 was relatively weak compared to that in yeast (Supplementary Fig. 9b). Such a discrepancy may be caused by either the mixed effect of the TM domains with and without SRP binding in the human dataset or the intrinsic mechanistic difference in cotranslational protein targeting mediated by SRP recognition between human and yeast.

## 3 Discussion

In this study, we proposed a deep learning based framework to predict ribosome stalling by integrating the underlying sequence features. To our best knowledge, our work is the first attempt to exploit the deep learning technique to predict ribosome stalling and model translation elongation dynamics based on the large-scale ribosome profiling data.

Our results demonstrated that ROSE is an accurate and robust method. We showed that although trained on a single dataset for each organism, ROSE can still accurately predict ribosome stalling events in several different datasets generated from other studies. More importantly, when predicting ribosome stalling events in regions not included in the training dataset, e.g., ramp regions and genes with low sequencing coverage, ROSE still showed comparable predition performance, indicating its potential ability to address the insufficient coverage problem that is common in current ribosome profiling data. Moreover, the correlation between RSS and multiple widely recognized conclusions about elongation dynamics further demonstrated the physiological relevance of the ribosome stalling events predicted by ROSE on a genome-wide scale. Collectively, these results showed that ROSE is able to capture important intrinsic sequence features to predict ribosome stalling based on high-throughput ribosome profiling data.

Similar to many other high-throughput sequencing techniques, the current analysis of ribosome profiling data is also faced with several technical challenges, e.g., aligning reads across exon-exon junctions, ambiguous mapping and sequencing bias [9]. In this study, we relied on several widely accepted data preprocessing approaches, e.g., RNA-Seq Unified Mapper (RUM) [97] in the alignment of splicing junction reads in GWIPS-viz [35] and the stringent normalization procedure proposed in [19], to at least partially remove the bias caused by these problems. Together with the accurate and robust prediction performance of ROSE as well as the physiologically relevant phenomena it detected, it is unlikely that the prediction of ROSE will suffer from the technical bias problem.

Our current study is a demonstration of applying ROSE in several specific scenarios in which ribosome stalling has been known to lead to significant physiological consequences. We believe that our ROSE framework will offer more insights into other important translation-related phenomena with the incorporation of more ribosome profiling data in future.

## 4 Methods

### 4.1 Data preparation, preprocessing and normalization

All datasets in this study were downloaded from GWIPS-viz [35], in which abundant ribosome profiling data have been maintained and preprocessed as in other widely accepted pipelines [9]. Here, we mainly focused on eukaryotic cells, including human and yeast cells. Two datasets with large data amount and high genome coverage in GWIPS-viz were used to train our deep learning model, including the human dataset of lymphoblastoid cell lines (LCLs) (denoted by Battle15) [56] and the yeast dataset of *S. cerevisiae* strain 288C (denoted by Pop14) [25]. In addition, we used other five additional datasets for cross-study validation, including three human datasets from different cell cycle stages (i.e., G1, S and M phases) of HeLa cells [59] (denoted by Stumpf13 G1, S and M, respectively) and two yeast datasets of strain SK1 [60] and starin BY4741 [61] (denoted by Brar12 and Young15, respectively).

Here we applied the normalization method introduced in [19] to remove the technical and experimental biases from the ribosome profiling data. More specifically, after mapping the ribosome profiling and mRNA-seq reads to the reference genome, their codon-level reads were first scaled by the mean coverage level within each gene, which canceled out the coverage differences among genes. Next, the scaled ribosome profiling reads were divided by the scaled mRNA-seq reads in the corresponding locations to eliminate the shared biases between these two fractions. After that, a logarithm operation was further performed to yield the final normalized ribosome footprint density (Supplementary Figs. 2a and 2b). Since some protein-coding genes can be poorly sequenced due to the issue of sequencing depth and the influence of differential expression, which may introduce unexpected biases to our analysis, here those normalized ribosome footprint densities from genes with sequencing coverage (i.e., the number of codon sites with both non-zero ribosome profiling and mRNA-seq reads divided by the total number of codons in the gene) less than 60% were excluded from our training and test datasets. Note that such a coverage cutoff was also used in [98], in which robustness of this cutoff has been demonstrated in the analysis of ribosome profiling data.

To label samples for a binary classifier detecting ribosome stalling events, we first tested several labeling thresholds based on the standard deviation of the normalized ribosome footprint density distribution. In particular, we considered four possible thresholds, including *µ, µ* + *σ, µ* + 2*σ* and *µ* + 3*σ*, where *µ* and *σ* represented the mean and the standard deviation of the normalized footprint density distribution, respectively. For each possible choice of threshold, those codon positions with normalized densities beyond the threshold were labeled as the ribosome stalling positions (i.e., positive samples), while the remaining were regarded as the background (i.e., negative samples). Then we trained four preliminary CNN models based on the datasets derived from these four thresholds, respectively. We also constructed a separate validation dataset that contained an equal number of samples randomly selected from six bins, including (−∞, *µ* – 2*σ*), [*µ* – 2*σ, µ – σ*), [*µ – σ, µ*), [*µ, µ* + *σ*), [*µ* + *σ, µ* + 2*σ*) and [*µ* + 2*σ*, ∞). The basic principle of choosing the optimal threshold was the expectation that our model with the best threshold should yield predictions best correlated with their corresponding experimentally observed values in the independent validation dataset. We performed such a test for both human and yeast datasets, from which *µ* + 2*σ* was determined as our final threshold (Supplementary Fig. 3).

After the above operations, the determined threshold was used to label samples, i.e., the codon sites with normalized footprint densities beyond the threshold were labeled as positive (i.e., foreground) samples, while the same number of codon sites randomly chosen from the remaining were labeled as negative (i.e., background) samples, which results in 109,770 and 20,902 samples for Battle15 and Pop14, respectively. For each dataset, we randomly selected 90% of the samples as training data and the remaining 10% as test data. The final performance of our model was mainly reported based on the test data. Note that here we excluded all the reads of the ramp regions (i.e., the first 50 codons at the 5’ ends of coding sequences) from the training data.

### 4.2 Model design

A convolutional neural network (CNN) is a specific type of neural network in deep learning, which has been widely used in common data science fields, such as computer vision [99] and natural language processing [100]. In particular, CNNs have also been used to model biological sequence data, e.g., the predictions of protein-nucleotide binding [44, 50] and effects of noncoding variants [47]. Generally speaking, a CNN is comprised of multiple local motif detectors (i.e., convolution operators) that are invariant with certain transformations, such as translation and rotation, and subsampling (i.e., pooling operators) for dimension reduction and efficient training. To further increase the learning capacity of the network, many layers of these operators are often stacked together, and then followed by several fully-connected layers, and finally the output layer.

In our framework, we first encode the input codon sequence using the one-hot encoding technique [101], that is, the *m*th codon is encoded as a binary vector of length 64, in which the *m*th position is one while the others are zeros, after indexing all 64 codons. Then the encoded information is fed into one convolutional layer and one pooling layer to learn the hidden features. In the convolutional layer, several one-dimensional convolution operations are performed over the 64-channel input data, in which each channel corresponds to one dimension of the input vector, and the weight matrix (i.e., kernel) can be regarded as the position weight matrix (PWM). More specifically, given a codon sequence *s* = (*c*_1_,…, *c*_*n*_) and the corresponding one-hot representation *S*, where *n* stands for the input length (here *n* = 61 as we extend the codon site of interest on both sides by 30 codons) and *c*_*i*_ represents the *i*th codon in the sequence, the convolutional layer computes X = conv(*S*), i.e.,

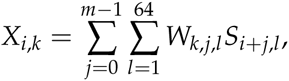

where 1 ≤ *i* ≤ *n − m* + 1,1 ≤ *k* ≤ *d,m* the kernel size, and *d* is the kernel number. Next, the rectified linear activation function (ReLU) is used to imitate the neuron activation, that is, the output of the convolutional layer is further processed by the activation function *Y* = ReLU(X), where

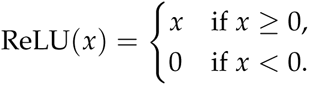

After convolution and rectification, we reduce the dimension of matrix *Y* using the max pooling operation, which computes the maximum value within a scanning window of size three and step size two. More specifically, given the upstream input *Y*, the max pooling operation computes Z = pool(*Y*), i.e.,

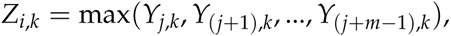

where *i* is the index of the output position, *j* is the index of the start input position, *k* is the index of the kernel, and *m* is the size of the scanning window during the pooling operation (here we choose *m* = 3).

To enable the local motif detectors to scan sequence motifs in different ranges synchronously, while not increasing the model complexity too much, here we propose a *parallel* architecture, which includes three kernels of different sizes, corresponding to short (5–7), mediate (8–9) and long (10–13) ranges, respectively. The outputs of these three kinds of convolution operators are further rectified and then subsampled independently and in parallel, and finally concatenated into a unified representation *U*. To calculate the final probability of a ribosome stalling event, the unified representation is directly fed to a sigmoid layer, which computes

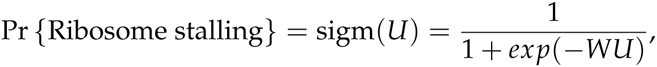

where *W* is the weight matrix of the sigmoid layer.

Note that the sequential (i.e., layer-wise) architecture in conventional CNNs, in which several convolutional and pooling layers are stacked together, can also detect motifs in different ranges. The reason that our parallel architecture can significantly reduce the model complexity comes from the fact that the parallelism simulates the SUM operation, e.g., (*a*_1_ + *a*_2_) + (*b*_1_ + *b*_2_), while the sequentiality mimics the PRODUCT operation, e.g., (*a*_1_ + *a*_2_) × (*b*_1_ + *b*_2_). Obviously, the computational complexity of the latter is much higher than that of the former. Our network reduction can be useful for relieving the potential overfitting problem during the training process. We note that a similar idea has also been proposed in [100]. However, the pooling operation in [100] is carried out over the whole convolutional layer without any window restriction, which is quit different from ours. In summary, a complete CNN in our deep learning framework can be formulated as

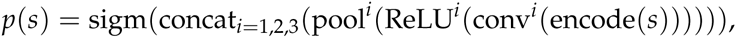

where *i* represents the kernel index in the parallel architecture, and encode(·), conv(·), ReLU(·), pool(·), concat(·) and sigm(·) represent the one-hot encoding, convolution, ReLU, max pooling, concatenation and sigmoid operations, respectively.

The above calculated probability *p*(*s*) is defined as the *intergenic ribosome stalling score* (also termed interRSS), which measures the likelihood of ribosome stalling at a codon position. To eliminate the interRSS bias among different genes, we further define the *intragenic ribosome stalling score* (also termed intraRSS) as follows,

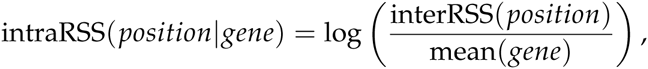

where interRSS(*position*) represents the interRSS of the codon position of interest and mean(*gene*) stands for the mean interRSS of the corresponding gene. When computing mean(*gene*), we exclude those codon positions in the ramp regions (i.e., the first 50 codons at the 5’ ends of coding sequences).

### 4.3 Model training and model selection

Given the training samples {(*s*_*i*_, *y*_*i*_)}*i*, the loss function of our model is defined as the sum of the negative log likelihoods (NLLs), i.e.,

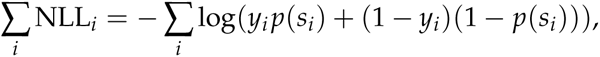

where *s*_*i*_ is the input codon sequence and *y*_*i*_ is the true label. To train the CNN, the standard batch gradient descent method with the error backpropagation algorithm is performed [53]. To further optimize the training procedure, we also apply several training strategies, including the mini-batch and momentum techniques [54]. In addition, we use the Adam algorithm for stochastic optimization to achieve an adaptive moment estimation [102]. To further overcome the overfitting issue, we also apply several regularization techniques, including *L*2-regularization-based weight decay [54], dropout [55] and early stopping [54].

The network structure and the aforementioned optimization techniques introduce a number of hyperparameters to our framework, such as the kernel size, kernel number, base learning rate, weight decay coefficient and the max number of training iterations. It is important to perform proper hyperparameter calibration and model selection for accurate modeling. Although we can achieve this goal using the conventional cross-validation strategies, it is generally time-consuming to test all possible combinations of these hyperparameters. To conquer this difficulty, here we propose a *one-way model selection* strategy for automatic and efficient hyperparameter calibration. In this strategy, we first arbitrarily choose the initial values of the hyperparameters from a candidate set. Then, we separate the hyperparameters into two groups, including those describing the network structure (denoted by *H*_1_), such as the kernel size and the kernel number, and those describing the optimization procedure (denoted by *H*_2_), such as the base learning rate and the weight decay coefficient. Next, by fixing the values of the hyperparameters in *H*_2_, we calibrate those hyperparameters in *H*_1_ using a three-fold cross-validation (CV) procedure, and determine their optimal values that achieve the best CV performance. Similarly, the hyperparameters in *H*_2_ are also calibrated via the three-fold CV procedure after fixing the previously determined values of the hyperparameters in *H*_1_. The final values of all hyperparameters of ROSE are provided in Supplementary Table 4. The ROC curves and AUROC scores of the CNNs with calibrated hyperparameters are shown in Supplementary Figs. 10a and 10b for the Battle15 and Pop14 datasets, respectively. Though we can carry out this procedure for more iterations (i.e., multi-way), our test results show that the one-way implementation generally yields satisfying prediction performance in this study.

After hyperparameter calibration and model selection, we train the final ROSE model using the whole training dataset. Due to the nature of non-convex optimization, random weight initialization may affect the search result of the gradient descent algorithm. Here, we use the Xavier initialization algorithm to automatically determine the initial scales of weights according to the number of input and output neurons [103]. To account for the potential initialization bias and further boost the prediction performance, we also implement an ensemble version of ROSE (termed eROSE), in which 64 CNNs are trained independently and then combined together to compute the final prediction score, i.e.,

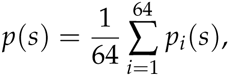

in which *p*_*i*_(*s*) represents the probability calculated by the ith CNN.

Our implementation of ROSE depends on the Caffe library [104], and the Tesla K20c GPUs are used to speed up the training process.

### 4.4 Statistical analysis on associations between diverse putative factors and ribosome stalling

Diverse factors, such as tRNA adaptation and mRNA secondary structure, can interplay with each other to affect the ribosome stalling tendency. To investigate whether a factor potentially correlates with ribosome stalling, we first identified those codon sites along the genome that were enriched with this factor, and then checked whether the predicted (intra)RSSes of these positions were significantly different from those of the background. In particular, given a factor, such as tRNA adaptation, we first computed its quantity (e.g., tAI) across the genome and then chose those codon sites whose quantities were in the top *N* list (*N* was set to 10,000 in our study). After that, we ran the Wilcoxon rank sum test to compare the (intra)RSSes of the chosen sites to those of a background dataset, which was generated by randomly selecting 10,000 ribosome occupancy sites from the genome. If the (intra)RSSes of the codon sites enriched with the factor and the background were significantly different, we said this factor correlates with ribosome stalling. In addition, we probed the correlations between different factors based on the background dataset, and found little correlation between these factors that we were interested in, except cAI, %MinMax and tAI (Supplementary Table 1). This enhanced our conclusions about the direct associations between diverse factors and ribosome stalling.

## Acknowledgments

This work was supported in part by the National Basic Research Program of China Grant 2011CBA00300, 2011CBA00301, the National Natural Science Foundation of China Grant 61033001, 61361136003 and 61472205, and China’s Youth 1000-Talent Program, the Beijing Advanced Innovation Center for Structural Biology. The authors are grateful to Drs. Q. Zhang and W. Chen for their helpful discussions about this work. They thank Ms. T. Chu for her help on preparing the figures in this paper.

## Author contributions

S.Z., H.H., J.Zhou and J.Zeng conceived the research project. J.Zeng supervised the research project. S.Z. preprocessed raw data, designed and implemented ROSE, and carried out model training and validation tasks. X.H. prepared the sequencing data of m^6^A modification. S.Z., H.H., J.Zhou, T.J. and J.Zeng performed the computational and statistical analyses. S.Z., H.H. and J.Zeng wrote the manuscript. All the authors discussed the test results and commented on the manuscript.

